# Automated cot-side tracking of functional brain age in preterm infants

**DOI:** 10.1101/848218

**Authors:** Nathan J. Stevenson, Lisa Oberdorfer, Maria-Luisa Tataranno, Michael Breakspear, Paul B. Colditz, Linda S. de Vries, Manon J. N. L. Benders, Katrin Klebermass-Schrehof, Sampsa Vanhatalo, James A. Roberts

## Abstract

**Objective:** A major challenge in the care of preterm infants is the early identification of compromised neurological development. While several measures are routinely used to track anatomical growth, there is a striking lack of reliable and objective tools for tracking maturation of early brain function; a cornerstone of lifelong neurological health. We present a cot-side method for measuring the functional maturity of the newborn brain based on routinely-available neurological monitoring with electroencephalography (EEG).

**Methods:** We used a dataset of 177 EEG recordings from 65 preterm infants to train a multivariable prediction of functional brain age (FBA) from EEG. The FBA was validated on an independent set of 99 EEG recordings from 42 preterm infants. The difference between FBA and postmenstrual age (PMA) was evaluated as a predictor for neurodevelopmental outcome.

**Results:** The FBA correlated strongly with the PMA of an infant, with a median prediction error of less than 1 week. Moreover, individual babies follow well-defined individual trajectories. The accuracy of the FBA applied to the validation set was statistically equivalent to the training set accuracy. In a subgroup of infants with repeated EEG recordings, a persistently negative predicted age difference was associated with poor neurodevelopmental outcome.

**Interpretation:** The FBA enables the tracking of functional neurodevelopment in preterm infants. This establishes proof of principle for growth charts for brain function, a new tool to assist clinical management and identify infants who will benefit most from early intervention.

## Introduction

Preterm birth is a substantial risk to infant health. While mortality rates have dropped considerably over recent years due to improvements in clinical care, these infants remain at significant risk of neurodevelopmental delay and a host of other chronic impairments in later life.^1,2^ It is therefore of critical importance to reduce the exposure of the preterm infant to neurological adversities while in the neonatal intensive care unit (NICU), and to identify those infants who will benefit most from early intervention.^3^ Recent advances in neurological care have stressed the need for improving early functional biomarkers of neurodevelopment to expedite cycles within clinical intervention trials.^4^

Monitoring physiological and anatomical growth is crucial for clinicians when optimizing the care of very or extremely preterm infants. Critical time periods for the direction of care are usually the first days after birth, the time of discharge from tertiary care to step-down units, as well as the follow-up visit at near term-equivalent age. Electroencephalography (EEG) is widely used for early therapeutic decisions and the prediction of neurodevelopmental outcome in preterm infants.^5–6^ Assessing brain maturity via the visual interpretation of the EEG during an infant’s stay in the NICU has been a part of clinical practice for decades.^7^ Its use complements traditional anatomical measures such as weight, length, and head circumference. However, the clinical use of EEG in the NICU is complicated by difficulties in interpretation and the availability of expertise to perform interpretation.^8^ Computer-assisted analysis presents an opportunity to solve both problems by providing simplified EEG measures that can be interpreted by clinical staff, on demand and in real time.

The concept of brain age is one such measure than can be automated.^9^ A lag between estimated functional brain age (FBA) from the EEG and the postmenstrual age (PMA) of the individual – the predicted age difference (PAD) – holds potential as a functional biomarker for use in neuro-intensive care. We have previously shown that it is possible to construct computational measures that emulate visually-observed features of maturation^10,11^ and correlate with pathological changes in neurological function.^12,13^ Recent advances in computational neuroscience suggest that the EEG contains markers of brain function that are not readily discernible by visual EEG review.^14^ Key information lies within the widespread network of intermittent bursting that dominates early cortical activity.^15^ This developmentally unique activity is known to be crucial for supporting neuronal growth and guiding early brain wiring.^16^ It changes rapidly over the last trimester, is sensitive to endogenous and exogenous disturbances, and is predictive of future neurodevelopment.^17,18, 19^ Here, we incorporated novel measures of the EEG into an analysis of the functional maturity of the preterm brain. We then determined the efficacy and validity of automated EEG analysis as a reliable biomarker of the functional maturity of the preterm brain and assessed its potential as a predictor of neurodevelopmental outcome.

## Materials and Methods

This study employed two different datasets of serial EEG recordings of preterm infants recorded from NICUs in different countries. The first dataset (recorded in Vienna: 65 infants, 177 EEG recordings) was used to train and evaluate the FBA measure, as well as investigate the use of FBA as a predictor of neurodevelopmental outcome (Fig 1). The second dataset (recorded in Utrecht: 42 infants, 99 EEG recordings) was used to validate the FBA measure trained on the first dataset. Infants were born before 29 weeks gestation, with EEG recorded serially at 25-39 weeks PMA (Vienna) or 25-34 weeks PMA (Utrecht). We used machine learning techniques to form an estimate of FBA using quantitative EEG (qEEG) variables that can be grouped into three categories: phenomenological analysis, burst analysis, and other recently-developed analyses (Table S1).

**Figure 1:**
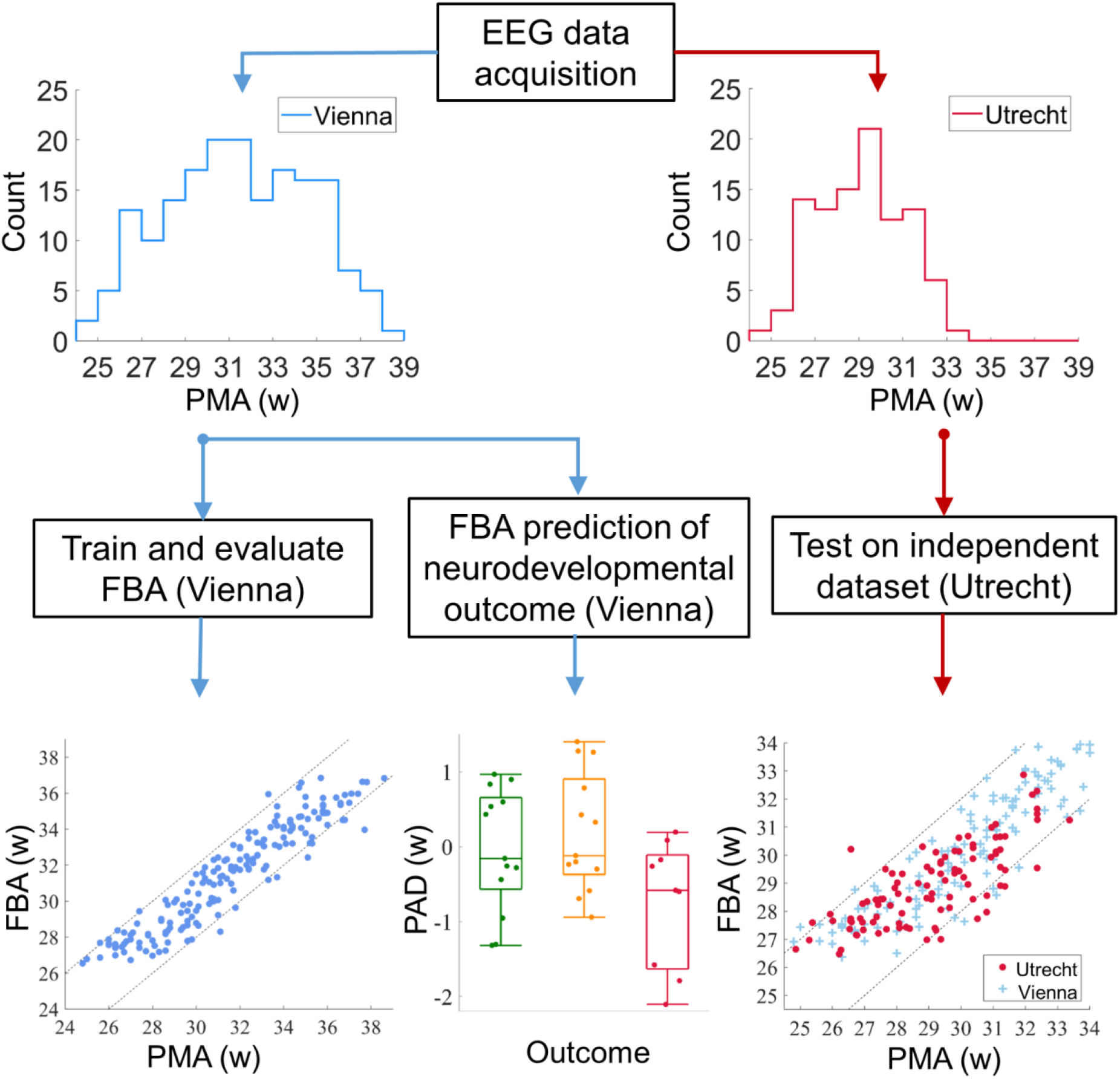
Data acquisition, training, evaluation and testing of the FBA. The histograms depict the distribution of EEG recordings with PMA (in weeks) in each dataset. The bottom row illustrates the analyses corresponding to each dataset. PAD is the predicted age difference between functional brain age (FBA) and post-menstrual age (PMA).

### Data Acquisition

The training dataset consisted of 67 preterm infants admitted to the NICU at the Vienna General University Hospital, Austria (see Table 1). Infants were included in the study cohort if they were born before 29 weeks gestational age, medically stable at the time of EEG recordings, and parental consent was received. EEG was acquired with nine scalp electrodes using a Brain Quick / ICU EEG (MicroMed, Treviso, Italy) at a sampling frequency of 256 Hz. Electrode positions reflect the 10-20 international system (modified for neonates) and were located at Fp1, Fp2, C3, C4, T3, T4, O1, O2, with a reference at Cz. A bipolar montage (double banana) was used in analysis: Fp1-C3, C3-O1, Fp1-T3, T3-O1, Fp2-C4, C4-O2, Fp2-T4, T4-O2. The EEG was recorded as soon as possible after birth and at fortnightly intervals until term equivalent age, where possible. Comorbidities and medications at the time of recording are listed in Table 1.

**Table 1:** A summary of infants and EEG recordings before and after application of artefact rejection. For outcome, percentages refer to the number of infants/recordings/1 h epochs that were in the initial set that passed the artefact rejection stage, for medications percentages are based on the number of recordings in the initial and post-artefact rejection sets, respectively and other values are presented as median (interquartile range). For intraventricular hemorrhage and periventricular leukomalacia, roman numerals indicate increasing grades of severity; assessed by cranial ultrasound.^20 a^ data missing from two infants.

| **Development: Vienna** | initial | post-artefact rejection |  |
| --- | --- | --- | --- |
| gestational age (weeks) | 25.3 (24.5-27.0) | 25.3 (24.5-27.0) |  |
| birthweight (g) | 707 (605-920) | 704 (604-922) |  |
| sex (m:f) <sup>a</sup> | 34:31 | 33:30 |  |
| PMA of EEG recording |  |  |  |
| 1<sup>st</sup> | 27.0 (26.6-29.4; n=67) | 27.9 (26.7-29.6; n=52) |  |
| 2<sup>nd</sup> | 30.8 (29.2-31.8; n=59) | 31.0 (29.5-31.8; n=43) |  |
| 3<sup>rd</sup> | 33.7 (32.0-34.4; n=54) | 33.6 (32.0-34.4; n=46) |  |
| 4<sup>th</sup> | 35.3 (33.1-36.4; n=37) | 35.2 (34.1-36.0; n=36) |  |
| 5<sup>th</sup> | 36.5 (35.2-37.4; n=16) | 36.4 (34.9-36.7; n=9) |  |
| 6<sup>th</sup> | 38.6 (n=1) | 38.6 (n=1) |  |
| intraventricular hemorrhage | 14 (I/II=10, III/IV=4) | 14 (I/II=10, III/IV=4) |  |
| periventricular leukomalacia | 2 (I/II=2) | 1 (I/II=1) |  |
| necrotizing enterocolitis | 3 | 3 |  |
| chronic lung disease | 19 | 19 |  |
| patent ductus arteriosus | 49 | 48 |  |
| medications at EEG recording |  |  |  |
| no medication | 15 (6%) | 10 (6%) |  |
| caffeine | 195 (83%) | 152 (86%) |  |
| morphine | 12 (5%) | 10 (6%) |  |
| inotropes | 5 (2%) | 5 (3%) |  |
| doxapram | 8 (3%) | 5 (3%) |  |
| anticonvulsants | 2 (1%) | 0 (0%) |  |
| dexamethasone | 1 (0.4%) | 1 (0.6%) |  |
| missing data | 14 (6%) | 9 (5%) |  |
|  | infants | Recordings | 1 h epochs |
| initial | 67 | 234 | 1686 |
| post-EEG assessment | 65 | 177 | 1137 |
| outcome |  |  |  |
| normal | 20 (100%) | 57 (76%) | 376 (65%) |
| mildly abnormal | 18 (95%) | 57 (78%) | 338 (73%) |
| abnormal | 16 (94%) | 41 (75%) | 238 (62%) |
| unknown | 11 (100%) | 22 (71%) | 185 (71%) |
| **Validation: Utrecht** | initial | post-EEG assessment |  |
| gestational age (weeks) | 26.9 (26.0-27.6) | 26.9 (26.1-27.6) |  |
| birthweight (g) | 920 (830-1068) | 920 (830-1070) |  |
| sex (m:f) | 27:16 | 26:16 |  |
| infants | 43 | 42 |  |
| recordings | 105 | 99 |  |
| 1 h epochs | 6561 | 6101 |  |

Each EEG recording was split into 1 h epochs (with a 75% overlap). Epochs with excessive artefact were excluded from further analysis (see Table 1). GA was defined according to the last menstrual period (LMP). If the LMP-based assessment of gestation deviated considerably from ultrasound findings in the first trimester, ultrasound measurements were used as a surrogate measure of LMP. PMA, defined as the sum of GA and postnatal age, was used as the benchmark, true age of an infant.

The neurodevelopmental outcome of infants was assessed at ages 1 y and 2 y (Bayley Scales of Infant Neurodevelopment II and III - BII and BIII, respectively; assessed against German norms). Of the 67 preterm infants initially included in the study, 2 were assessed with BII at 1 y, 19 were assessed with BII at 2 y, 35 were assessed with BIII at 2 y, and 11 infants were not assessed for neurodevelopmental outcome. Infants were stratified according to outcome using the following rules applied to the latest available assessment (either 1 y or 2 y): normal (mental developmental index and physical developmental index > 85 [scale II]; or cognitive index, language index, and motor index > 85 [scale III]), or abnormal (either mental developmental index or physical developmental index <70 [scale II]; or either cognitive index, language index, or motor index < 70 [scale III]). Those infants with intermediate scores that did not fit into the normal or abnormal categories were categorized as mildly abnormal.

A summary of the database before and after the rejection of epochs with an excessive amount of artefact is presented in Table 1. Data collection was approved by the local ethics committee and written, informed, parental consent was received for each infant included in the database (Medical University Vienna, Austria; study protocol EK Nr 67/2008).

### Data Acquisition for Independent, Validation Dataset

We used an independent dataset to validate the single and multivariable models of age prediction. This validation dataset contained EEG recordings from 43 neonates admitted to the NICU at the Wilhelmina Children’s Hospital, Utrecht, Netherlands. The data were collected as part of a multi-center European study.^21^ Infants were included in this study if they were born less than 28 weeks GA and informed, written parental consent was received. Infants were excluded if the presence of chromosomal or congenital abnormalities were identified and if the neuro-monitoring was performed with devices other than the BrainZ BRM3 monitor (Natus Medical Incorporated, Seattle, USA). Long duration EEG recordings (~72 h) were recorded as close as possible to admission, followed by shorter recordings (~4-6 h) at weekly intervals (up to a post-natal age of approximately four weeks). EEG was recorded with a BrainZ BRM3 monitor and needle electrodes at a sampling frequency of 256 Hz. Two derivations were recorded and used in the analysis: F4-P4 and F3-P3. All neonates had a neurological examination and psychological testing at 30 months of corrected age (Bayley Scales of Infant and Toddler Development III; assessed against Dutch norms). Neonates with normal neurodevelopmental outcome at this age were included in the validation cohort (normal was defined as per the Vienna dataset; *n* = 43 infants satisfied these criteria). The data collection protocol was approved by the local medical ethics committee.

### EEG preprocessing

EEG recorded in an intensive care environment is prone to contamination from electrical activity that is not cortical in origin. All data were filtered with a high pass filter (Butterworth, 4^th^ order, cutoff frequency at 0.5 Hz) and then a low-pass filter (Butterworth, 6^th^ order, cutoff frequency at 16 Hz) to eliminate high frequency activity that is more commonly associated with artefacts, including muscle activity.^8^ EEG recordings were then segmented into 1 h epochs. To account for further artefacts, yet include as many 1 h EEG epochs as possible, we used automated rejection of EEG epochs with excessive artefact. We did not analyze epochs with considerable spatial imbalance in amplitude (EEG derivations with a factor of 2 difference in mean amplitude from any other derivation), persistent excessive amplitude (greater than 25% of the recording with burst amplitudes greater than 500 μV), or persistent low amplitude (greater than 50% of the recording with EEG activity less than 5 μV).

### Prediction of Post-Menstrual Age using qEEG

Single and multivariable models of PMA were calculated using regression analysis. These models used a combination of summary measures of the EEG (qEEG variables), calculated on 1 h epochs, as an input and generated a prediction of PMA as an output. The qEEG variables used in this study can be grouped into three categories: phenomenological analysis (*m* = 46), burst analysis (*m* = 40), and other analysis paradigms (*m* = 10). Phenomenological analysis extracts qEEG variables that mirror the visual interpretation of the EEG.^10–11^ Burst analysis extracts qEEG variables that identify important characteristics of highly irregular (“crackling”) noise, through analysis of EEG bursts.^14^ Other advanced analyses extract qEEG variables that represent complex characteristics of the preterm EEG such as entropy, global connectivity, and cross-channel coupling.^22–28^

Leave-one-subject-out cross-validation was used when generating models with a single or multivariable input and FBA as output. In the case of *N* subjects (infants), a training set consisting of the qEEG variables from *N*-1 subjects was used to train the regression model that outputs a FBA. The accuracy of the FBA was then calculated using the left-out subject. The process was repeated until all subjects had been left out, allowing accuracy to be estimated on the entire cohort. Single multivariable model parameters were estimated using support vector regression with a medium Gaussian kernel (a kernel scale of 10, a box constraint equal to the interquartile range of PMA/1.349 and epsilon equal to the box constraint/10). Support vector regression is tolerant of redundant and irrelevant variables; nonetheless, we implemented a process of variable selection to rank the importance of each qEEG variable to the determination of age and to reduce the computational burden of the multivariable model. Backwards selection was used (4-fold cross-validation within the training set), with the mean square error between FBA and PMA as a cost function to be minimized.

### Independent Validation

A multivariable model was trained on all available data in the Vienna dataset and then applied to the independent dataset collected at the Wilhelmina Children’s Hospital, Utrecht, Netherlands. The range of PMA and electrode recording configurations were not identical between the EEG recordings from Vienna and Utrecht. To overcome this heterogeneity, the multivariable model was trained on fronto-central derivations (Fp1-C3, Fp2-C4) in the Vienna dataset and applied to the fronto-parietal derivations in the Utrecht dataset. This was one of the closest approximations, in terms of position and distance between electrodes, given the available configurations of the Vienna dataset (we also tested centro-temporal derivations). Model efficacy was only compared across a similar PMA range between the two datasets (24-33 weeks PMA).

### Statistical Analysis

A prediction was made on a 1 h epoch of EEG; if multiple EEG epochs exist per recording then the average predicted age per recording was used. The goodness-of-fit between predicted age (single and multivariable models) and PMA was evaluated using the correlation coefficient (Pearson’s). The bias, variance, and absolute error between predicted age and PMA were also used as measures of goodness-of-fit (*13*). The use of repeated (serial) measures allowed the application of a linear mixed effects model (LMM) where the model output was a fixed effect and the infant ID was a random effect. The adjusted *r* value was used to assess the goodness-of-fit taking into account multiple recordings from each infant. Statistical comparisons of measures of the goodness-of-fit between EEG metrics for the prediction of PMA were performed using resampling methods (bootstrap). A correlation coefficient was deemed significantly different if the 95% confidence interval of differences (estimated via a bootstrap) did not span zero, i.e., was either positive or negative. Differences in prediction accuracy (absolute error) between the Vienna (training) and Utrecht (validation) sets were evaluated using equivalence testing with a two, one-sided t-tests (TOST) procedure based on Welch’s t-test.^29^ We used a mean difference in absolute error of ±0.5 weeks as a conservative equivalence boundary based on the results of previous work which reported an absolute prediction error of approximately ±1 week.^11^ Differences in FBA trajectories (FBA subtracted from PMA averaged across all recordings per infant) between outcome groups were tested using a one-way ANOVA, with Levene’s test for homogeneity of group variances and a post hoc analysis performed using Tukey’s Range test to correct for multiple comparisons. FBA trajectories in each group were assessed to determine if they were significantly different from zero using a t-test. For post-hoc analyses, Cohen’s D statistic, with small sample size correction, was used to estimate the effect size between groups. All tests were two-tailed and used a level of significance of 0.05.

## Results

### PMA prediction using a single variable FBA

Across all metrics tested, the qEEG variable that had the highest correlation with PMA was the asymmetry of average burst shape (Fig 2A), which exhibits a strong linear relationship with bursts becoming more symmetric with increasing PMA (Fig 2B). Several additional qEEG variables were strongly associated with PMA (Table S1). Metrics that were not reliably predictive of PMA were varied in nature and included several relative band powers and measures of burst duration.

**Figure 2.**
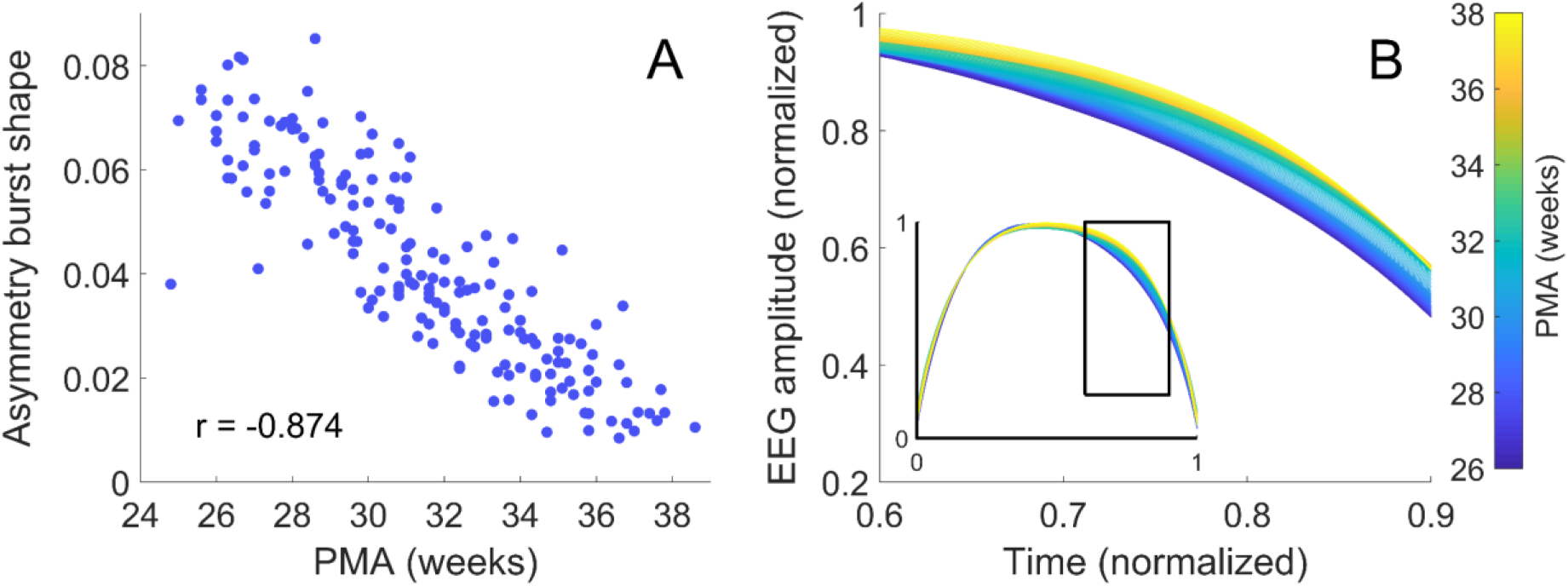
Changes in burst characteristics with post-menstrual age (PMA). (**A**) Asymmetry of average burst shape versus PMA (*r* is Pearson’s linear correlation coefficient). (**B**) Average burst shape of the EEG amplitude grouped according to PMA with fortnightly steps from 25 weeks; the inset shows the entire average burst. The changes seen in (B) are best represented by measures of burst asymmetry.

### PMA prediction using the multivariable FBA

Combining several qEEG variables into a multivariable model improves the prediction accuracy of the FBA (Table 2). Assessed within a leave-one-out cross-validation, the multivariable FBA model had a significantly higher correlation with PMA than a single variable model based on the single best variable (asymmetry of the burst shape) for models based on bursts, phenomenological, and other newly proposed qEEG variables (Δ*r* = 0.109, 95% CI: 0.059 to 0.162; Δ*r* = 0.095, 95% CI: 0.045 to 0.150; Δ*r* = 0.094, 95% CI: 0.057 to 0.142; *n* = 177, respectively). The multivariable model using burst qEEG variables also had a significantly higher correlation with PMA than multivariable models based on phenomenological or other newly proposed qEEG variables (Δ*r* = 0.030, 95% CI: 0.001 to 0.062; Δ*r* = 0.027, 95% CI: 0.005 to 0.049; *n* = 177, respectively).

**Table 2:** The performance of several multivariable FBA models for predicting PMA in preterm infants on training (cross-validation) and validation datasets. *r* is the correlation coefficient, *n* is the number of recordings included in analysis, *m* is the number of qEEG variables used in the model (for variable selection this is the median number across folds of the cross-validation), 95% CI is the 95^th^ percentile of the confidence interval, IQR is inter-quartile range, FC and CO denote a FBA trained on fronto-central and centro-occipital derivations, respectively.

| | $r$ [95%CI] | bias<br>(weeks) | variance<br>(weeks) | absolute<br>difference<br>(weeks)<br>[IQR] | $\pm 1$<br>week<br>(%) | $\pm 2$<br>weeks<br>(%) |
| --- | --- | --- | --- | --- | --- | --- |
| Phenomenological ($n = 177$; $m = 46$) | 0.894<br>[0.859-0.919] | -0.1 | 2.1 | 0.9<br>[0.4-1.6] | 55 | 83 |
| Other<br>($n = 177$; $m = 10$) | 0.896<br>[0.866-0.920] | -0.1 | 2.1 | 0.9<br>[0.5-1.5] | 53 | 84 |
| Bursts<br>($n = 177$; $m = 40$) | 0.923<br>[0.905-0.940] | -0.2 | 1.6 | 0.9<br>[0.3-1.4] | 60 | 89 |
| Variable Selection ($n = 177$; $m = 53$) | 0.938<br>[0.922-0.952] | -0.1 | 1.3 | 0.7<br>[0.4-1.3] | 63 | 90 |
| **Validation:** Vienna<br>($n = 134$; $m = 53$) | 0.900<br>[0.873-0.929] | -0.2 | 1.1 | 0.6<br>[0.3-1.2] | 64 | 92 |
| **Validation:** Utrecht<br>(FC: $n = 99$; $m = 53$) | 0.765<br>[0.665-0.846] | 0.1 | 1.5 | 0.9<br>[0.4-1.3] | 61 | 90 |
| **Validation:** Utrecht<br>(CO: $n = 99$; $m = 53$) | 0.674<br>[0.554-0.758] | -0.3 | 2.0 | 1.0<br>[0.4-1.6] | 53 | 80 |

Incorporating qEEG variables into a multivariable model, via a variable selection procedure, further improved the accuracy of the FBA (Table 2). The FBA estimator identified PMA to within 2 weeks for 90% of recordings, with a median absolute error of 0.7 weeks. A scatter plot of FBA versus PMA exhibits a clear linear trend (Fig 3A), with a tight clustering of FBA within ±2 weeks of the PMA. The performance of this FBA, which contained a mixture of burst, phenomenological, and other recently-developed EEG variables, was significantly higher than multivariable models based on only phenomenological analysis, burst analysis or other analyses alone (Δ*r* = 0.045, 95%CI: 0.020 to 0.073; Δ*r* = 0.015, 95%CI: 0.002 to 0.028; and Δ*r* = 0.042, 95%CI: 0.024 to 0.063, respectively; *n* = 177). Variable selection resulted in a median of 53 variables (IQR; 49-55; *n* = 65 folds, see Table S1).

**Figure 3:**
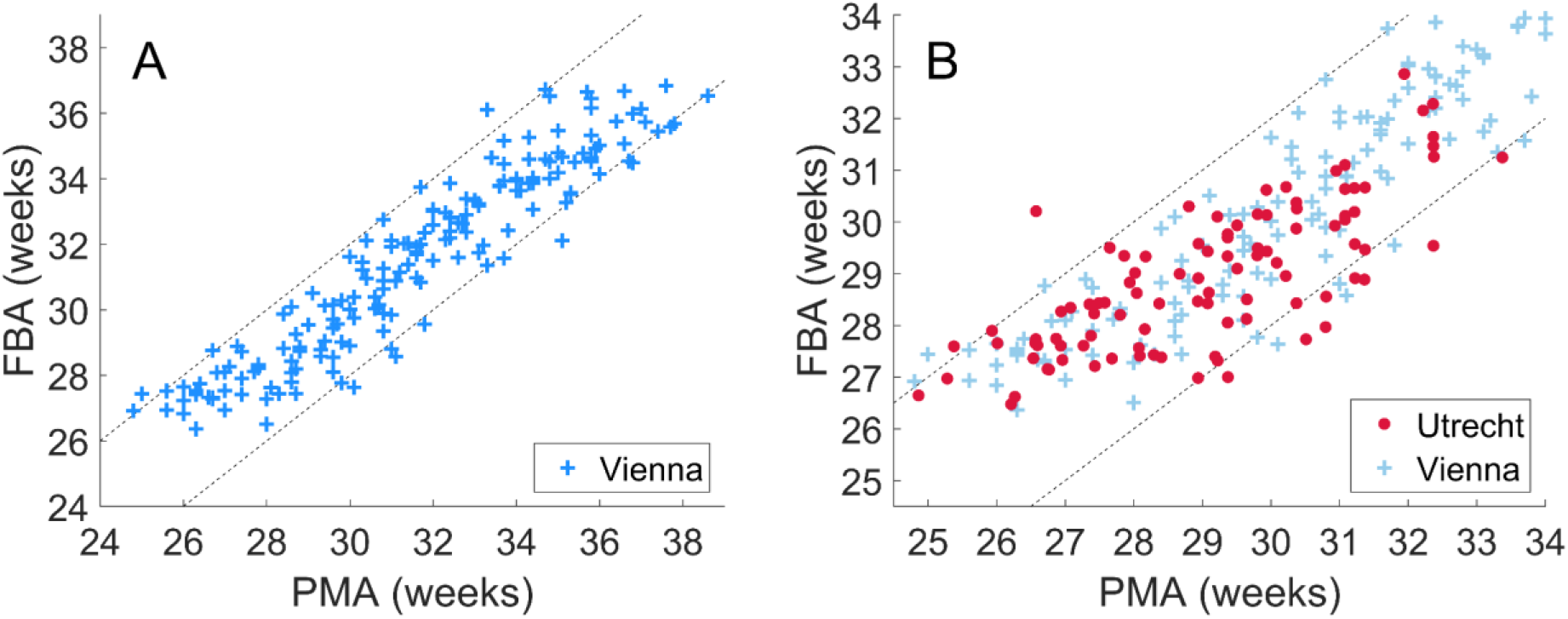
The correlation between a multivariable FBA and PMA. (**A**) The multivariable FBA, with variable selection, evaluated on the Vienna dataset via leave-one-subject-out cross-validation over the full range of EEG recording PMAs (24-38 weeks). (**B**) The multivariable FBA trained on the Vienna dataset and applied to an independent dataset recorded from Utrecht over the full range of EEG recording PMAs from the Utrecht dataset (24-34 weeks). Dashed lines denote ±2 weeks difference between FBA and PMA.

### Validation of FBA on an independent dataset

To directly address the generalizability of our results, we validated the multivariable model in an independent dataset. We found that the multivariable model trained on Vienna data (evaluated via cross-validation) and applied to the Utrecht dataset performs near physiological limits in prediction accuracy, with 90% of epochs correctly identified to within ±2 weeks (Table 2). The absolute error between the FBA and PMA (accuracy), when applied to the Utrecht data, was equivalent to the cross-validation results from the Vienna dataset across a similar range of PMA (Fig 3B: *p* < 0.001; TOST, equivalence boundary of ±0.5 weeks). Training with centro-temporal recordings also resulted in a FBA that was strongly correlated with PMA when applied to the Utrecht dataset (Table 2).

### FBA for tracking individual growth and predicting neurodevelopmental outcomes

The accuracy of FBA in tracking cot-side development raises the idea that FBA may be useful for individualized assessment of functional maturity. We used linear mixed modelling to account for serial recordings from individual infants which resulted in an adjusted correlation of *r* = 0.978 (95%CI: 0.974-0.987; *n* = 65). The improvement in correlation over a point-wise estimate implies that individual infant trajectories are more highly correlated with PMA than the cohort average. In other words, infants tend to follow their individual growth trajectories (Fig 4A), and the FBA is able to track these trajectories with high accuracy. In a subgroup of infants with more than two serial recordings (a median PMA range of 6.2 weeks, IQR: 4.6 to 7.5 weeks; Fig 4B), the average predicted age difference (PAD: difference between FBA and PMA) was significantly associated with neurodevelopmental outcome (One-way ANOVA: F statistic = 3.980, df = 2, *p* = 0.029; *n* = 35, 3 groups: normal, mildly abnormal, abnormal – see Fig 4C; group variances were homogenous; Levene’s Test: *p* = 0.82). Infants with abnormal outcome (*n* = 9) had a PAD that was significantly less than infants with mildly abnormal outcome (*n* = 13) (Cohen’s D = 1.12, *p* = 0.025, corrected for multiple comparisons using Tukey’s Range Test). The estimated PAD in these infants was also significantly below 0 weeks (t-test: Cohen’s D = 0.661, *p* = 0.035; *n* = 9) suggesting a persistent delay in brain maturation (i.e., negative PAD) in infants with abnormal outcome. These differences were not apparent when including infants with less than three serial EEG recordings (ANOVA: F statistic = 0.0112, df = 2, *p* = 0.894; *n* = 54), suggesting that multiple recordings may be required to assess a PAD associated with neurodevelopmental outcome.

**Figure 4:**
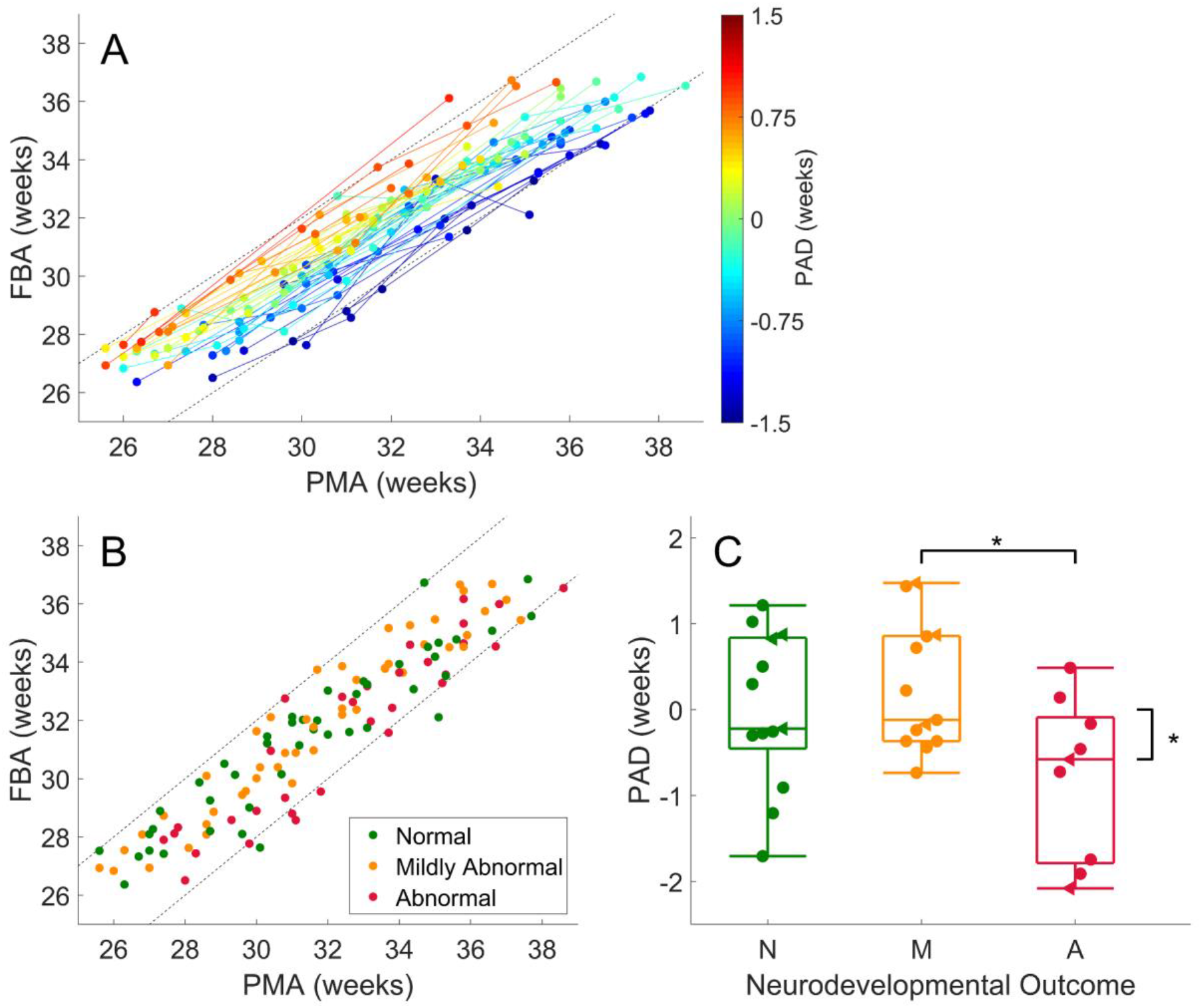
Functional brain age prediction using a multivariable model of quantitative EEG measures. (**A**) Maturational trajectories of individual infants, with at least two serial recordings per infant; *n* = 54, colored according to the average differences between FBA and PMA (PAD: predicted age difference) in each infant. The color bar denotes the PAD in weeks. (**B**) Scatter plot of the subgroup of data, with at least three serial recordings, used to evaluate the prediction error for outcome prediction; *n* = 35, colored according to neurodevelopmental outcome. Straight dashed lines denote a difference of plus or minus 2 weeks between PMA and predicted age. (**C**) Subgroup analysis of EEG predicted age minus PMA with respect to outcome was graded as N – normal (minimum Bayley’s score > 85), M – mildly abnormal (minimum Bayley’s score between 70 and 85) and A – abnormal (minimum Bayley’s score < 70). The asterisks denote *p* < 0.05 between outcome groups and when testing each outcome group against a null hypothesis of zero mean EEG maturity. Data points in have been shifted for clarity of presentation and are denoted with filled circles. Data points represented by triangles are infants with intra-ventricular hemorrhage.

## Discussion

The brain matures rapidly in early life with a wide range of structural and functional indices changing over time spans as short as a few weeks. Here, we showed that automated analysis of preterm EEG can be used to track maturation of cortical function with high accuracy. This analysis converts the EEG into an ‘age’ trend (FBA) that can be considered as a biomarker of maturity in preterm EEG. This multivariable prediction of age from the EEG enables the estimation of functional brain maturity to within 1-2 weeks of PMA; an accuracy that generalized to an independent validation dataset acquired under a considerably different EEG recording environment. The margin of error is far lower than similar predictions in preterm infants based on functional neuroimaging with fMRI and orders of magnitude lower than what is achieved over later stages of life using EEG or MRI (error margins of 5-10 years).^30, 31, 32^ Our findings are also comparable to an array of somatic anatomical methods over similar preterm age ranges based on measures of femur length, head circumference, weight, and structural MRI (cortical folding, thickness).^33, 34^ This supports the concept of rapid and distinct changes in anatomy and physiology throughout the preterm period and suggests that physiological and anatomical growth are strongly intertwined.^18, 35, 36^

The multivariable model developed here advances previous work that was designed to capture key visual elements of EEG review for age prediction. Incorporating burst measures based on the analysis of crackling noise resulted in the most accurate single variable model, improved multivariable model accuracy, and provided a potential framework to explain the mechanistic origins of rapidly evolving preterm EEG signals. The existence of asymmetric burst shapes replicates our previous findings in independent datasets when identifying pathological changes in the EEG at or near birth.^13, 17^ We also validated several recently proposed qEEG variables of maturation: suppression curve, mPLI, global ASI, multi-scale entropy, and path length (coherence) as excellent predictors of age prediction in the preterm period. This supports the use of automated measures of EEG (qEEG) for the extraction of useful information in excess of visual interpretation.

We also successfully validated the model's robustness on unseen data. This showed that the prediction accuracy of the multivariable model holds when translating to a dataset collected within a different clinical environment and with different recording parameters (e.g. amplifier, electrode type, number, and location). This is a crucial hurdle for the clinical translation of new methods, which is impossible to establish in a dataset acquired under uniform conditions.

Notably, the independent validation dataset was collected using a 4-channel recording montage that is commonly used in brain monitoring with the amplitude integrated EEG.^37^ This validation on heterogeneous data establishes the wider clinical applicability of determining functional brain age from EEG.

Various measures of growth are commonly used in health care. The finding that neurological dysfunction manifests as immaturity in the EEG is intuitively appealing, and indeed, has been a cornerstone of clinical EEG review for decades.^38–40^ This hypothesis can only be accurately tested with qEEG measures that are strongly correlated with age. We show that most phenomenological measures used in clinical EEG research, such as inter-burst interval, EEG amplitude, or spectral power, are only weakly correlated with age, which challenges their applicability for maturational EEG assessment. More recently proposed qEEG measures (such as asymmetry and sharpness of burst shape, suppression curve, mPLI, MSE, and path length) and multivariable models of age are strongly correlated with PMA and, therefore, more relevant for maturational analyses.

We show that infants follow individual functional maturation trajectories in a highly predictable manner. Analysis of these trajectories with measures such as PAD (the difference between FBA and PMA) can, potentially, be used to predict neurodevelopmental outcome. This suggests that early neurological adversities become embedded in cortical function (EEG recordings).^41^ The potential of PAD measures was only apparent in infants with multiple recordings over a wide range of PMA within the preterm period. Summarizing across multiple recordings resulted in a PAD that was able to capture both acute and chronic delays in functional maturation.^5^

Limited sample sizes, a reality when studying critically ill infants, mean that the reported links between PAD and outcome cannot take into account the variety of factors that may confound this result such as physiological challenges, routine cares, and interventions experienced by preterm infants during their stay in intensive care. These factors will also confound the FBA. There is evidence to suggest that several of these challenges, interventions (of which, many are designed to accelerate maturation), and other relevant clinical variables can influence EEG activity and, therefore, contribute to variability in the FBA and, therefore, PAD. These variables include the difference between GA and PMA (intra-uterine vs extra-uterine maturation), post-natal adaption, medications, ventilation, birthweight, kangaroo care/infant massage, and gender.^42–47^ We aim to investigate these effects in future work to differentiate them from other potential causes of inter-subject variability such as natural variability in the course of in-utero growth.^48^ Nevertheless, FBA provides an accurate prediction of PMA even when based on a small training sample with an array of potentially confounding factors that were not explicitly modelled. This is a reassuring finding, but the incorporation of EEG trends such as FBA into routine clinical practice requires an FBA trained on a larger cohorts and independent validation with other well characterized clinical cohorts to tease out the role of potential confounders, and determine the effect of acute, chronic or even longer-term changes in the underlying EEG.

The clinical potential of FBA is twofold. First, tracking of individual growth trajectories is becoming an important part of individualized medicine for preterm infants.^49, 50^ Tracking FBA provides a crucial functional complement to anatomical growth charts, being sensitive to the functional consequence of perinatal adversities specific to early neurological development. Second, these analyses may have an important role in clinical trials, as recent progress in early therapeutic interventions has been hampered by delays due to the assessment of outcome several years after birth. The use of very early measures of neurodevelopment, like FBA, could lead to dramatically expedited study cycles by allowing more dynamic, adaptive study designs with optimized sample sizes and research questions.^51^ The estimation of FBA also has clear applications in developmental neuroscience, where the assessment of maturation based on cortical function can be used to benchmark models of early human neurological development across species and within human brain organoids.^52, 53^

## Acknowledgments

This work was supported by the National Health and Medical Research Council of Australia (JAR, MB, SV, PBC, APP1144936; MB, APP1118153), the Rebecca L. Cooper Foundation (JAR, PG2018109), Lastentautiensäätiö (SV), the Finnish Academy (SV, 313242, 288220), Fonds zur Förderung der Wissenschaftlichen Forschung (KKS, FWFKLI237) and the European Commission (LdV, MJNLB; LSHM-CT-2006-036534).

## Author contributions

NJS, SV and JAR contributed to study conceptualization; NJS, JAR and SV undertook preliminary investigations, NJS performed the formal analysis; LO, MLT, LdV, KKS, SV contributed to data collection and curation; JAR, MB, SV, and PBC acquired funding for the study; and all authors contributed to the writing of the paper.

