## Supplementary Table S1 for "Automated cot-side tracking of functional brain age in preterm infants"

**Table S1:** Correlation between single variable prediction based on a single qEEG variables and post-menstrual age (PMA) within a leave-one-subject-out cross-validation. Type defines: B - a set representing measures of bursts, P - a set representing phenomenological interpretation of the EEG, O - a set representing other measures in the literature, [1] denotes bursts defined according to [1], [2] denote bursts defined according to [2], and selected denotes variable selection frequency.

| rank | qEEG variable | type | $r$ | selected |
|---|---|---|---|---|
| 1 | Asymmetry (all) | B | 0.874 | 29 |
| 2 | Asymmetry (500ms-1s) | B | 0.851 | 64 |
| 3 | Sharpness (500ms-1s) | B | 0.845 | 51 |
| 4 | Burst Number[1] (4-8s) | B | 0.843 | 45 |
| 5 | Suppression Curve | O | 0.842 | 63 |
| 6 | Burst Number[1] (2-4s) | B | 0.842 | 65 |
| 7 | Asymmetry (250-500ms) | B | 0.829 | 35 |
| 8 | mean Phase Locking Index | O | 0.822 | 62 |
| 9 | Path length (coherence) | O | 0.811 | 65 |
| 10 | Burst Number[1] (1-2s) | B | 0.807 | 29 |
| 11 | Multi-Scale Entropy ($\Delta$ 5 scales) | O | 0.807 | 64 |
| 12 | Multi-Scale Entropy (mean) | O | 0.794 | 65 |
| 13 | Multi-Scale Entropy (maximum) | O | 0.778 | 60 |
| 14 | Mean Burst Duration[1] | B | 0.777 | 24 |
| 15 | Burst Number[1] (125-250ms) | B | 0.772 | 64 |
| 16 | Sharpness (1-2s) | B | 0.720 | 57 |
| 17 | Burst Number[1] (500ms-1s) | B | 0.713 | 28 |
| 18 | Asymmetry (1s-2s) | B | 0.676 | 28 |
| 19 | Slope of linear fit to burst area vs burst duration[1] | B | 0.667 | 63 |
| 20 | Number of bursts[1] | B | 0.656 | 65 |
| 21 | Alpha (truncated power law fit to CDF of burst size) | B | 0.644 | 53 |
| 22 | Intercept of linear fit to asymmetry vs burst duration[1] | B | 0.643 | 35 |
| 23 | Sharpness (62.5-125ms) | B | 0.630 | 65 |
| 24 | Activity Synchrony Index (global) | O | 0.627 | 1 |
| 25 | Sharpness (1-2s) | B | 0.618 | 49 |
| 26 | Range EEG (5th percentile, quiet sleep) | P | 0.595 | 2 |
| 27 | Range EEG (5th percentile, full recording) | P | 0.592 | 47 |
| 28 | Cross-Channel Correlation (full recording) | P | 0.572 | 58 |
| 29 | Inter-Burst Interval[2] (95th percentile, full recording) | P | 0.572 | 13 |
| 30 | Inter-Burst Interval[2] (median, quiet sleep) | P | 0.567 | 1 |
| 31 | Cross-Channel Correlation (quiet sleep) | P | 0.566 | 24 |
| 32 | Alpha (truncated power law fit to CDF of burst duration) | B | 0.565 | 12 |
| 33 | Sharpness (s) | B | 0.563 | 7 |
| 34 | Inter-Burst Interval[2] (median, full recording) | P | 0.553 | 65 |
| 35 | Bursts per hour[2] (full recording) | P | 0.535 | 28 |
| 36 | RMS Inter-Burst Interval[2] (full recording) | P | 0.530 | 5 |
| 37 | Bursts per hour[2] (quiet sleep) | P | 0.527 | 6 |
| 38 | Inter-Burst Interval[2] (95th percentile, quiet sleep) | P | 0.525 | 3 |
| 39 | Range EEG (95th percentile, full recording) | P | 0.524 | 54 |
| 40 | RMS Inter-Burst Interval[2] (quiet sleep) | P | 0.522 | 1 |
| 41 | Lambda (truncated power law fit to CDF of burst size) | B | 0.521 | 19 |
| 42 | Sharpness (125-250ms) | B | 0.505 | 45 |
| 43 | Intercept of linear fit to burst size vs burst duration[1] | B | 0.502 | 65 |
| 44 | EEG envelope (95th percentile, full recording) | P | 0.495 | 57 |
| 45 | Relative Alpha Power (quiet sleep) | P | 0.481 | 65 |
| 46 | Burst Number[1] (62.5-125ms) | B | 0.480 | 37 |
| 47 | EEG envelope (5th percentile, quiet sleep) | P | 0.478 | 1 |
| 48 | Activity Synchrony Index (hemispheric) | O | 0.474 | 28 |
| 49 | Log-Likelihood Ratio of Fit (truncated power law fit to CDF of burst size) | B | 0.473 | 8 |
| 50 | Sharpness (all) | B | 0.467 | 59 |
| 51 | Range EEG (95th percentile, quiet sleep) | P | 0.460 | 42 |
| 52 | Relative alpha power (full recording) | P | 0.440 | 63 |

| | | | | |
|---|---|---|---|---|
| 53 | Inter-Burst Interval[2] (5th percentile, full recording) | P | 0.440 | 29 |
| 54 | Inter-Burst Interval[2] (5th percentile, quiet sleep) | P | 0.433 | 10 |
| 55 | Total spectral power (full recording) | P | 0.433 | 26 |
| 56 | EEG envelope (5th percentile, full recording) | P | 0.425 | 33 |
| 57 | Temporal Theta Power (full recording) | P | 0.425 | 44 |
| 58 | EEG Envelope (95th percentile, quiet sleep) | P | 0.416 | 16 |
| 59 | Asymmetry (2-4s) | B | 0.353 | 24 |
| 60 | Log-Likelihood Ratio of Fit (truncated power law fit to CDF of burst duration) | B | 0.341 | 60 |
| 61 | Total spectral power (quiet sleep) | P | 0.338 | 35 |
| 62 | Burst duration[2] (median, quiet sleep) | P | 0.337 | 17 |
| 63 | Temporal Theta Power (quiet sleep) | P | 0.328 | 47 |
| 64 | Asymmetry (125-250ms) | B | 0.32 | 29 |
| 65 | Burst duration (5th percentile, quiet sleep) | P | 0.324 | 33 |
| 66 | $s_{min}$ (truncated power law fit to CDF of burst duration) | B | 0.323 | 15 |
| 67 | Burst duration[2] (5th percentile, full recording) | P | 0.304 | 16 |
| 68 | Activity Synchrony Index (quiet sleep) | P | 0.283 | 0 |
| 69 | Segment Rate | O | 0.246 | 36 |
| 70 | Activity Synchrony Index (full recording) | P | 0.245 | 65 |
| 71 | Coefficient of variation (burst durations)[1] | B | 0.231 | 61 |
| 72 | Slope of linear fit to sharpness vs burst duration[1] | B | 0.227 | 19 |
| 73 | Burst duration[2] (median, full recording) | P | 0.227 | 3 |
| 74 | Relative delta 1 power (quiet sleep) | P | 0.221 | 58 |
| 75 | Relative delta 2 power (quiet sleep) | P | 0.210 | 23 |
| 76 | Range EEG (median, quiet sleep) | P | 0.206 | 14 |
| 77 | Lambda (truncated power law fit to CDF of burst duration) | B | 0.202 | 45 |
| 78 | Burst duration[2] (95th percentile, full recording) | P | 0.193 | 10 |
| 79 | EEG Envelope (median, quiet sleep) | P | 0.187 | 2 |
| 80 | RMS burst duration[2] (full recording) | P | 0.169 | 2 |
| 81 | Relative delta 2 power (full recording) | P | 0.166 | 65 |
| 82 | Relative delta 1 power (full recording) | P | 0.151 | 60 |
| 83 | Asymmetry (4-8s) | B | 0.130 | 14 |
| 84 | Slope of linear fit to asymmetry vs burst duration[1] | B | 0.120 | 6 |
| 85 | Relative theta power (full recording) | P | 0.084 | 56 |
| 86 | Asymmetry (62.5-125ms) | B | 0.068 | 42 |
| 87 | Range EEG (median, full recording) | P | 0.062 | 49 |
| 88 | Sharpness (4-8s) | B | 0.058 | 0 |
| 89 | EEG Envelope (median, full recording) | P | 0.043 | 65 |
| 90 | $s_{min}$ (truncated power law fit to CDF of burst area) | B | -0.013 | 47 |
| 91 | Relative theta power (quiet sleep) | P | -0.024 | 62 |
| 92 | Range EEG (bandwidth) | O | -0.088 | 30 |
| 93 | Burst duration[2] (95th percentile, quiet sleep) | P | -0.106 | 0 |
| 94 | RMS burst duration[2] (quiet sleep) | P | -0.167 | 0 |
| 95 | Burst number[1] (250-500ms) | B | -0.221 | 40 |
| 96 | Intercept of linear fit to sharpness vs burst duration[1] | B | -0.310 | 4 |
